# A novel chimeric coronavirus spike vaccine combining SARS-CoV-2 RBD and scaffold domains from HKU-1 elicits potent neutralising antibody responses

**DOI:** 10.1101/2025.07.16.665240

**Authors:** Veronica Zoest, Wen Shi Lee, Lydia Murdiyarso, Lauren Burmas, Phillip Pymm, Robyn Esterbauer, Andrew Kelly, Hannah G Kelly, Isaac Barber-Axthelm, James P Cooney, Kathryn C Davidson, Merle Dayton, Courtney E McAleese, Marianne Gillard, Karen Hughes, Martina L Jones, Marc Pellegrini, Wai-Hong Tham, Ben Hughes, Stephen J Kent, Adam K Wheatley, Jennifer A Juno, Hyon-Xhi Tan

## Abstract

The receptor binding domain (RBD) of the SARS-CoV-2 spike is the major target for neutralising antibodies elicited by current vaccines. Using small domains such as the RBD as vaccine immunogens, however, may constrain the availability of CD4 T follicular helper (TFH) cells and impact immunogenicity. We engineered a novel chimeric trimeric RBD (CTR) glycoprotein, replacing the RBD of human coronavirus HKU-1 spike with SARS-CoV-2 RBD of either ancestral (WT) or Omicron BA.2 strains. This strategy maintains a native trimeric conformation of the RBD, while providing additional sources of CD4 T cell help via the HKU-1 spike scaffold. In C57BL/6 mice, CTR-BA.2 prime-boost vaccination elicited high anti-BA.2-RBD IgG and neutralising titres, matching responses in animals immunised with native SARS-CoV-2 spike proteins. GC B cells elicited by CTR-BA.2 were predominantly WT^+^/BA.2^+^ cross- reactive, and TFH cells predominantly recognised HKU-1 epitopes, demonstrating scaffold-directed T cell help. Macaques prime-boost immunised with CTR-WT similarly elicited high anti-RBD IgG, anti-spike IgG and neutralising responses, comparable to native spike-vaccinated animals. In draining lymph nodes of CTR-WT vaccinated macaques, RBD-specific GC B cells were present at elevated levels. In contrast to the murine studies, lymph node-draining TFH responses in macaques were broadly elicited against RBD, NTD/S2 or HKU-1-derived peptides. Although native SARS-CoV-2 spike was also highly immunogenic in animal models, our findings establish the chimeric glycoprotein design as a strategy to overcome the poor immunogenicity of the SARS-CoV-2 RBD by engaging CD4 TFH cells, while maintaining the ability to elicit protective neutralising responses.

**One sentence summary:** A chimeric glycoprotein design preserves SARS-CoV-2 RBD antigenic conformation enabling elicitation of neutralising responses, while allowing recruitment of HKU-1 scaffold-directed CD4 helper responses to support the humoral response.

## Introduction

The SARS-CoV-2 spike (S) glycoprotein is the primary immunogen in most COVID-19 vaccines and is highly effective at eliciting neutralising antibodies. Embedded within the spike protein, the receptor binding domain (RBD) mediates viral engagement with cellular angiotensin-converting enzyme 2 (ACE2) receptors and subsequent cell entry. While neutralising antibodies can target distal regions across the spike including the N-terminal domain (NTD)(*1*) and S2 domain(*2*), the majority of potent neutralising activity is associated with antibodies that directly block RBD-ACE2 interactions(*3–6*). Vaccines that preferentially focus antibody responses toward the RBD are therefore a potential pathway to maximising protective vaccine efficacy.

The RBD subunit itself can be used as a vaccine immunogen(*7*, *8*). However, we and others have demonstrated that monomeric RBD is intrinsically poorly immunogenic in pre-clinical models, due in part to suboptimal levels of CD4 T cell help (*9*, *10*). To augment the low immunogenicity of small protein subdomains while concurrently minimising repetitive boosting of non-neutralising epitopes, vaccine antigens can be embedded into scaffolds or carriers, which can provide T cell help via inclusion of heterologous sequences. Examples of this approach include the fusion of SARS-CoV- 2 NTD and RBD(*11*, *12*) and chimeric influenza hemagglutinin proteins comprised of an H5 globular head and H1 stem(*13*). Immunogenicity can be further improved through incorporation of trimerisation motifs that maintain the native-like trimeric conformation of type I glycoproteins and often augment humoral immune responses compared to monomeric antigens(*14*, *15*).

The spike proteins of endemic human coronaviruses (hCoVs) provide a potential path to present the SARS-CoV-2 RBD in a trimeric format with ample CD4 T cell help. In addition to ubiquitous seroconversion to hCoVs in children and adults(*16–18*), we and others have previously demonstrated hCoV-specific CD4 T cell memory responses are highly prevalent in the population(*19–21*). We therefore hypothesised that a chimeric immunogen could harness hCoV-specifc CD4 T cells, in this case derived from the distantly related beta-coronavirus HKU-1, to augment the intrinsically poor immunogenicity of the SARS-CoV-2 RBD, while limiting the biogenesis or recall of immunity to the SARS-CoV-2 NTD and S2 domains that have limited neutralising potency.

Here, we designed a stabilised chimeric trimeric RBD (CTR) glycoprotein, consisting of an HKU-1 backbone scaffold (NTD and S2 domains) with the native RBD substituted for the SARS-CoV-2 RBD. We characterised CTR immunogens in murine and macaque models, demonstrating the efficient elicitation of HKU-1-specific T helper responses and the induction of serum binding and neutralising antibody activity broadly comparable to that elicited with native full-length SARS-CoV-2 spike. This work demonstrates that a chimeric glycoprotein is a viable vaccine design strategy that combines trimerisation and robust CD4 T cell helper responses to drive neutralising responses to intrinsically poorly immunogenic glycoprotein subdomains.

## Results

### Chimeric spike glycoprotein displays human ACE2 binding, heterogenous HKU- 1 and SARS-CoV-2 RBD antigenicity, and a trimeric tertiary structure

Sequences encoding SARS-CoV-2 RBD from ancestral (WT) or BA.2 strains were engineered into an HKU-1 scaffold comprising the NTD and S2 domains, which we termed CTR-WT and CTR-BA.2 respectively (Fig 1A). CTR constructs were stabilised in a pre-fusion conformation via the inclusion of S2P modifications(*22*) and in a trimeric conformation via incorporation of a T4 fibritin fold-on motif(*23*). CTR proteins were expressed using mammalian cell culture and purified using affinity and size exclusion chromatography, with a trace profile similar to that of trimeric SARS-CoV-2 spike (Supplementary Figure S1).

**Figure 1.**
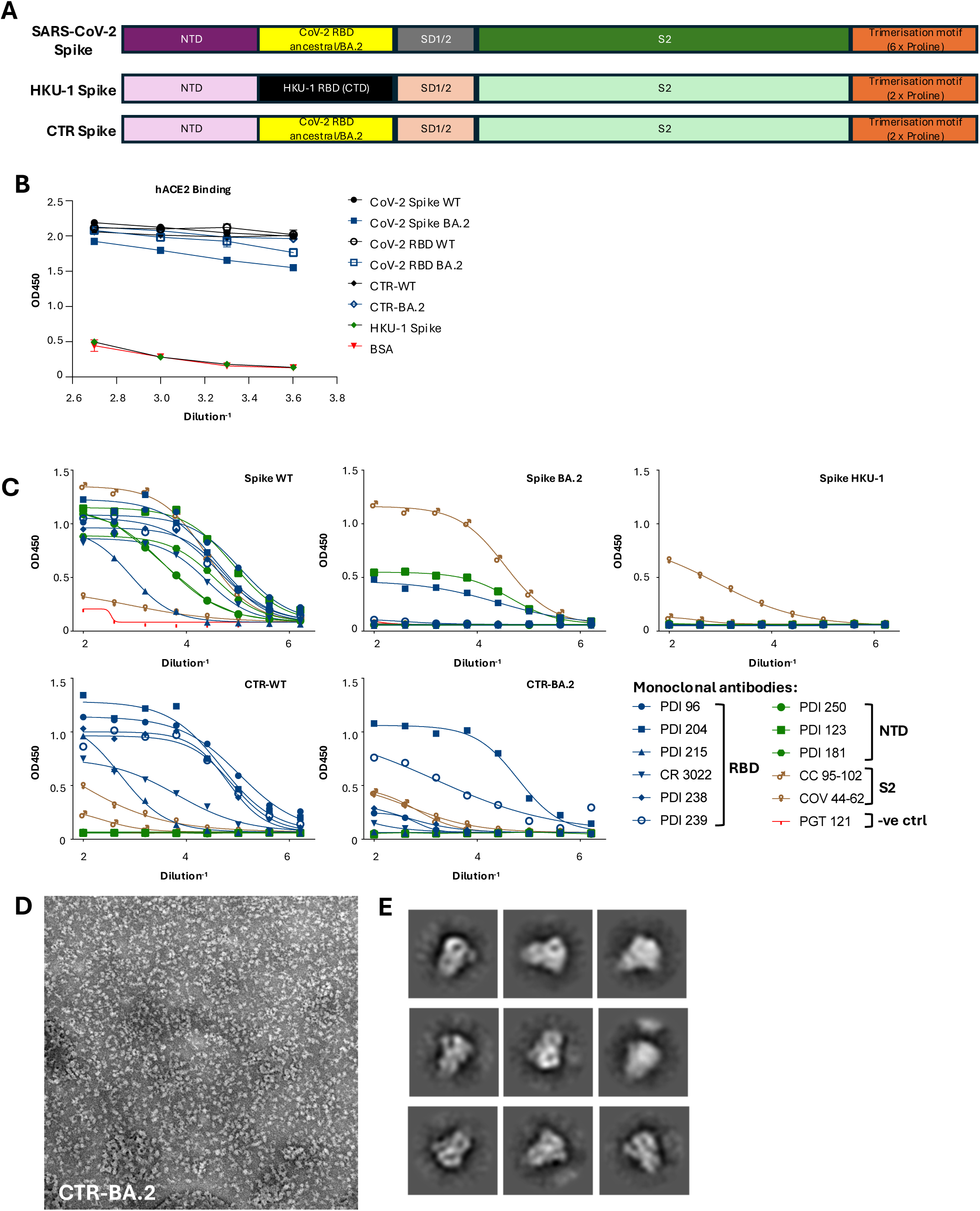
Design and characterisation of chimeric spike glycoproteins. (A) Comparison of SARS-CoV-2, HKU-1 and CTR spike protein constructs. CTR spike was engineered by replacing the RBD (CTD) of HKU-1 with SARS-CoV-2 RBD. (B) Human ACE2 binding capacity of native spike, RBD and CTR proteins. (C) Antigenicity of SARS-CoV-2, HKU-1 and CTR spike proteins determined by ELISA using a panel of monoclonal antibodies. (D) Negative stain electron micrograph of CTR BA.2, with (E) selected and magnified 2D class images highlighting trimer-like conformations.

We first performed a human ACE2 (hACE2) binding assay to validate the functional integrity of the RBD within these chimeric antigens (Fig 1B). Native spike or RBD proteins from WT or BA.2 strains displayed high binding activity to hACE2, with comparable binding observed for CTR-WT and CTR-BA.2 antigens. In contrast, binding was absent for the HKU-1 spike control, which is specific for TMPRSS2(*24*). We also utilised monoclonal antibodies (mAbs) with known specificities to SARS-CoV- 2 RBD, NTD, and S2 domains to antigenically characterise the spike proteins (Fig 1C). RBD-specific mAbs recognised both spike WT and CTR-WT proteins, confirming appropriate presentation of the RBD in the CTR protein. In contrast, NTD- and S2- specific mAbs bound spike WT but not CTR. Numerous mutations present in the BA.2 spike and RBD abrogated the binding of most mAbs, with the exception of a conserved S2 mAb that bound spike BA.2, and an RBD specific mAb (PDI-204) that bound both spike BA.2 and CTR-BA.2. As expected, the HKU-1 spike was recognised by a known human beta-coronavirus cross-reactive S2 mAb (COV44-62)(*25*, *26*).

The structure of CTR-BA.2 was further analysed using negative stain transmission electron microscopy (Fig 1D). Selected 2D class images revealed CTR-BA.2 in a trimeric conformation, with distinct S1 head and S2 stalk domains visible (Fig 1E). Overall, the chimeric coronavirus spike combining SARS-CoV-2 RBD and scaffold domains from HKU-1 displayed antigenic integrity and a quaternary structure suitable for further characterisation in immunogenicity studies.

### CTR is immunogenic in mice and elicits neutralising antibody responses comparable to native SARS-CoV-2 spike

To first establish that the chimeric spike glycoprotein elicits protective responses against SARS-CoV-2, BALB/c mice were prime-boost immunised with combinations of OVA control protein, spike WT, spike HKU-1 and/or CTR-WT co-formulated with Addavax (an MF-59-like squalene adjuvant) and subsequently challenged with mouse infectious SARS-CoV-2 N501Y virus (Supplementary Figure S2A). Three days post-challenge, lungs were excised and viral loads were measured. Animals vaccinated twice with control OVA protein displayed high titres of SARS-CoV-2 in the lung. In contrast, mice immunised with at least one dose of CTR-WT showed limited to no viral replication in lungs, comparable to protection seen in animals vaccinated with spike WT (Supplementary Figure S2B). Given the protective potential of the CTR platform, we performed subsequent in-depth immunogenicity studies in murine models using the CTR-BA.2 antigen, as Omicron BA.2 was the predominant circulating viral variant during the study period.

The immunogenicity of CTR-BA.2 was assessed in C57BL/6 mice by prime-boost vaccination and compared to animals immunised with native SARS-CoV-2 spike from the WT or BA.2 strains (Fig 2A). CTR-BA.2 was immunogenic, eliciting serum IgG titres against RBD WT or BA.2 at day 14 post-boost that were comparable to spike- immunised animals (Fig 2B). Serological responses elicited by the HKU-1 backbone were observed in CTR-BA.2 immunised animals, with lower, but detectible anti-HKU- 1 IgG titres seen in native spike-immunised animals presumably arising from cross- reactive S2 epitopes(*25*). Neutralising activity against WT live virus was elicited only in animals vaccinated with spike WT, while BA.2-specific neutralisation was comparable across all groups (Fig 2C). None of the antigens elicited neutralising antibodies against the much more distant XBB1.5 variant.

**Figure 2.**
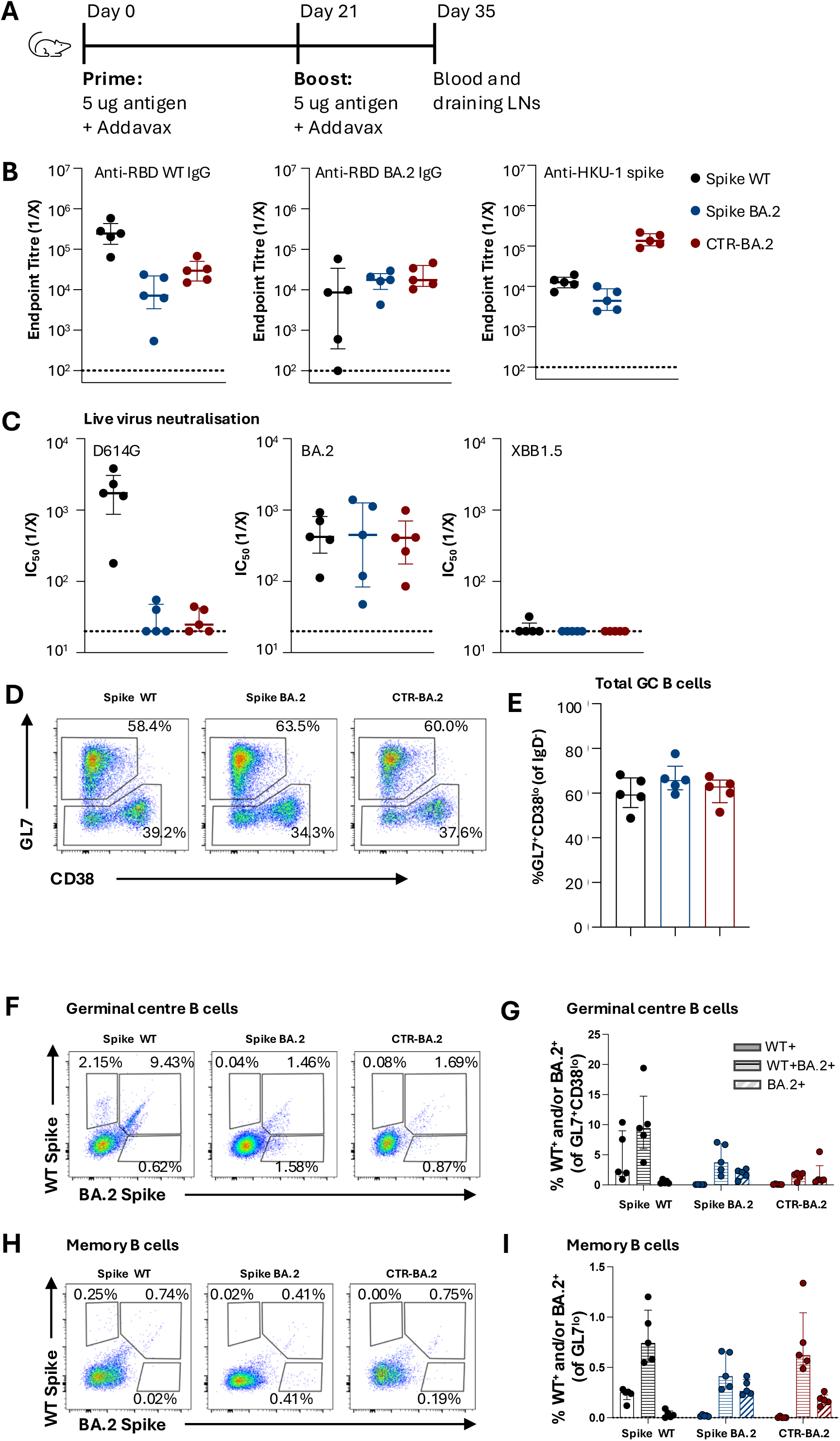
Prime-boost immunisation of CTR in C57BL/6 mice. (A) Immunisation schedule of mice vaccinated with SARS-CoV-2 WT spike, SARS-CoV-2 BA.2 spike or CTR BA.2 spike (n = 5 / group). (B) Serum endpoint titres of anti-RBD, -RBD BA.2 and -HKU-1 spike at day 35 determined by ELISA. (C) Neutralisation activity in serum against SARS-CoV-2 D614G, BA.2 and XBB1.5 assessed by live virus neutralisation assay. (C-D) Dotted lines denote the detection cut off. (D) Representative staining of GC B cells (GL7+ CD38lo) and memory B cells (GL7lo). (E) Frequency of total GC B cells. (F) Representative staining and (G) quantification of WT and/or BA.2 spike-specific GC B cells. (H) Representative staining and (I) frequency of WT and/or BA.2 spike-specific memory B cells.

To gain insight into the potential cellular event driving the serum neutralising responses observed, the elicitation of B cell responses in the draining iliac and inguinal lymph nodes (LN) were assessed by flow cytometry using recombinant spike WT or BA.2 probes (gating in Supplementary Fig S3A). CTR-BA.2 immunised animals exhibited frequencies of germinal centre (GC) B cells (GL7^+^ CD38^lo^) that were comparable to animals immunised with native spike (Fig 2D-E). In all vaccine groups, the majority of antigen-specific GC and memory (GL7^lo^) B cells were cross-reactive (WT^+^ BA.2^+^) (Fig 2F-I). Probe-specific GC B cell frequencies were lowest in the CTR- BA.2 group, most likely due to the presence of S2- and NTD-specific B cells in the native spike-immunised groups.

These data demonstrate that the placement of SARS-CoV-2 RBD onto a chimeric HKU-1 backbone is immunogenic and elicits serological and neutralising antibody responses comparable to native spike glycoproteins. Mirroring this neutralising antibody response, CTR-BA.2 antigen elicits GC and memory B cell responses in draining LNs of prime-boost immunised animals specific against SARS-CoV-2 spike.

### CTR elicits T helper responses predominantly from epitopes within the HKU-1 scaffold

In contrast to whole spike, we and others previously linked the poor immunogenicity of the SARS-CoV-2 RBD to low availability of CD4 helper epitopes in mouse models(*9*, *10*, *27*). We therefore immunised mice with a single dose of spike WT, spike HKU-1, CTR-WT or CTR-BA.2 and characterised the quantity and specificity of TFH (CD4^+^ CXCR5^++^ PD-1^++^ CD44^hi^) in the draining LN by measuring CD154 (CD40L) expression after restimulation with whole spike or RBD proteins (Fig 3A; gating in Supplementary Fig S3B). As previously shown(*9*), spike-specific TFH at 14 days post-vaccination in C57BL/6 mice are targeted to epitopes localised outside of the RBD (Fig 3B-C). In HKU-1-immunised animals, TFH were specific to epitopes unique to the HKU-1 spike and not cross-reactive with SARS-CoV-2 spike. In mice vaccinated with CTR-WT or CTR-BA.2, TFH predominantly recognised epitopes within the spike HKU-1 scaffold (S2 and NTD domains), although some reactivity was observed towards epitopes within the BA.2 RBD (Fig 3C). Similar responses were observed in the non-TFH memory CD4 T cell compartment (Supplementary Fig S3C).

**Figure 3.**
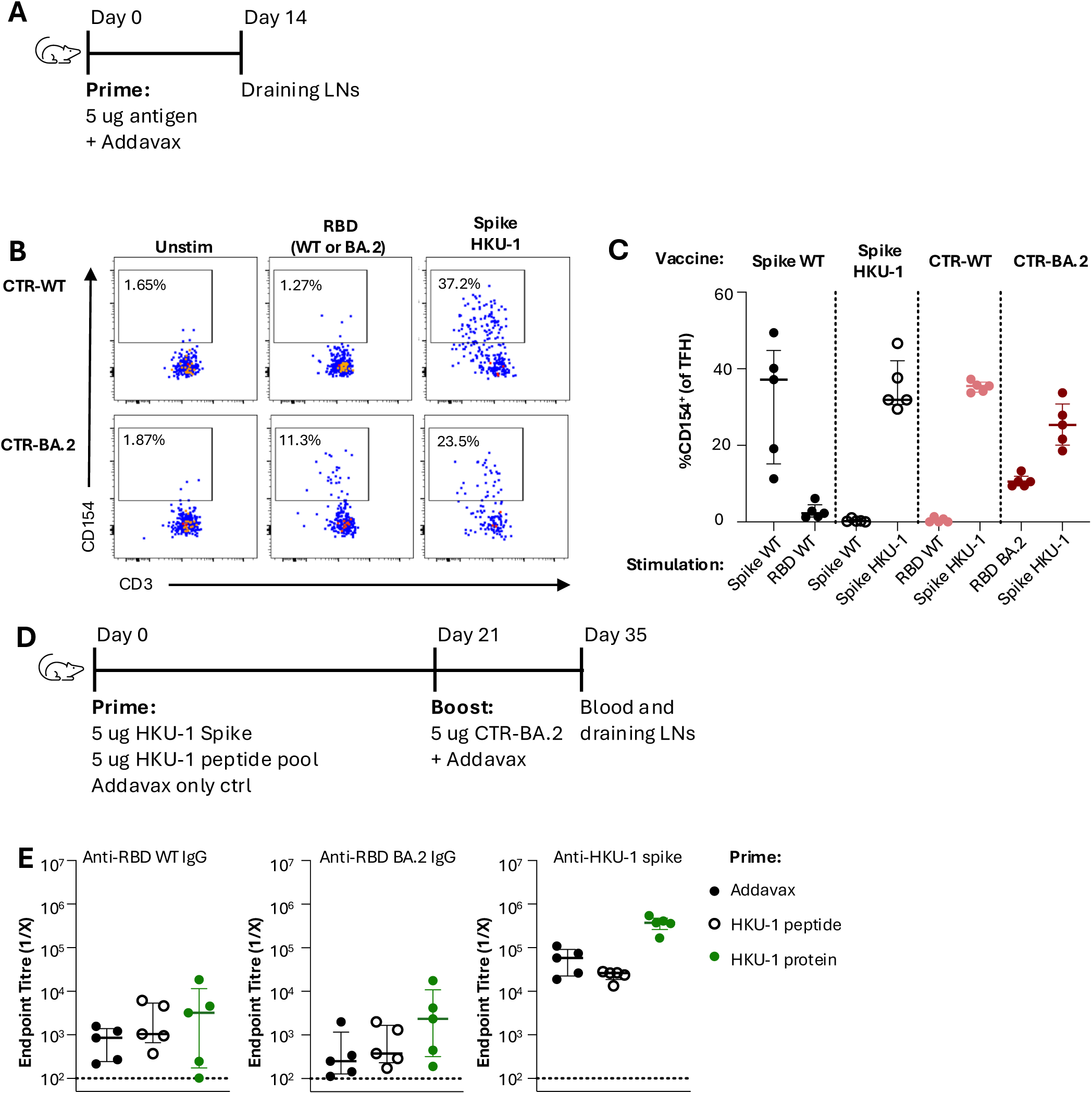
Assessing TFH responses in CTR-immunised C57BL/6 mice. (A) Immunisation schedule to assess primary TFH responses in mice vaccinated with SARS-CoV-2 WT spike, HKU-1 spike, CTR-WT, CTR-BA.2 spike (n = 5 / group). (B) Representative staining of lymph node draining antigen-specific TFH measured by expression of activation marker CD154. (C) Frequencies of antigen-specific CD154+ TFH cells upon restimulation with whole spike or RBD proteins. (D) Immunisation schedule to assess impact of pre-existing HKU-1 immunity on CTR immunogenicity. Mice were primed with either Addavax alone, a peptide pool spanning the HKU-1 spike to prime CD4 T cells or whole HKU-1 spike protein to prime antibody, memory B and CD4 T cells (n = 5 / group). (E) Serum endpoint titres of anti-RBD, -RBD BA.2 and -HKU-1 spike at day 35 determined by ELISA. Dotted lines denote the detection cut off.

Given the near-universal humoral and CD4 T cell memory to HKU-1 spike in adult populations(*21*), we next assessed whether pre-existing HKU-1 immunity would impact the immunogenicity of CTR-BA.2 (Fig 3D). Given a lack of a murine infection model for HKU-1 and to mimic HKU-1 immunity, mice were primed with either Addavax alone, a peptide pool spanning the HKU-1 spike (to prime CD4 T cells) or the HKU-1 spike protein (to prime antibody, memory B and CD4 T cells). HKU-1 priming had no impact on WT or BA.2 RBD IgG titres, although protein priming did drive higher titres of HKU-1 spike-specific IgG following the CTR-BA.2 boost (Fig 3E). Overall, these data demonstrate that the CTR antigen can establish robust GC responses through recruitment of HKU-1-specific TFH, even in the absence of high-quality RBD-specific CD4 help. Importantly, pre-existing HKU-1-specific CD4 T cells or serological responses do not appear to inhibit the immunogenicity of CTR antigens.

### CTR efficiently elicits germinal centres and recalls memory B cells in animals with prior spike immunity

Clinically approved SARS-CoV-2 booster vaccines are now administered in a complex immune landscape, including prior COVID-19 infection and/or primary vaccination. We therefore assessed the immunogenicity of CTR-BA.2 in animals with prior spike WT immunity (Fig 4A). Animals were immunised twice with spike WT and rested 8 weeks before prime-boost vaccination with spike WT, spike BA.2 or CTR-BA.2. Antibody titres against either RBD WT or RBD BA.2 were similarly boosted in all groups following both the 3^rd^ and 4^th^ dose (Fig 4B). As expected, animals receiving CTR-BA.2 exhibited significantly higher HKU-1 spike titres after the 4^th^ dose, due to repeated exposure to the majority of the HKU-1 spike. Neutralising responses at day 105 were similar across all groups against WT or BA.2 viruses, while responses to XBB1.5, a more antigenically distant Omicron strain, were low or absent (Fig 4C).

**Figure 4.**
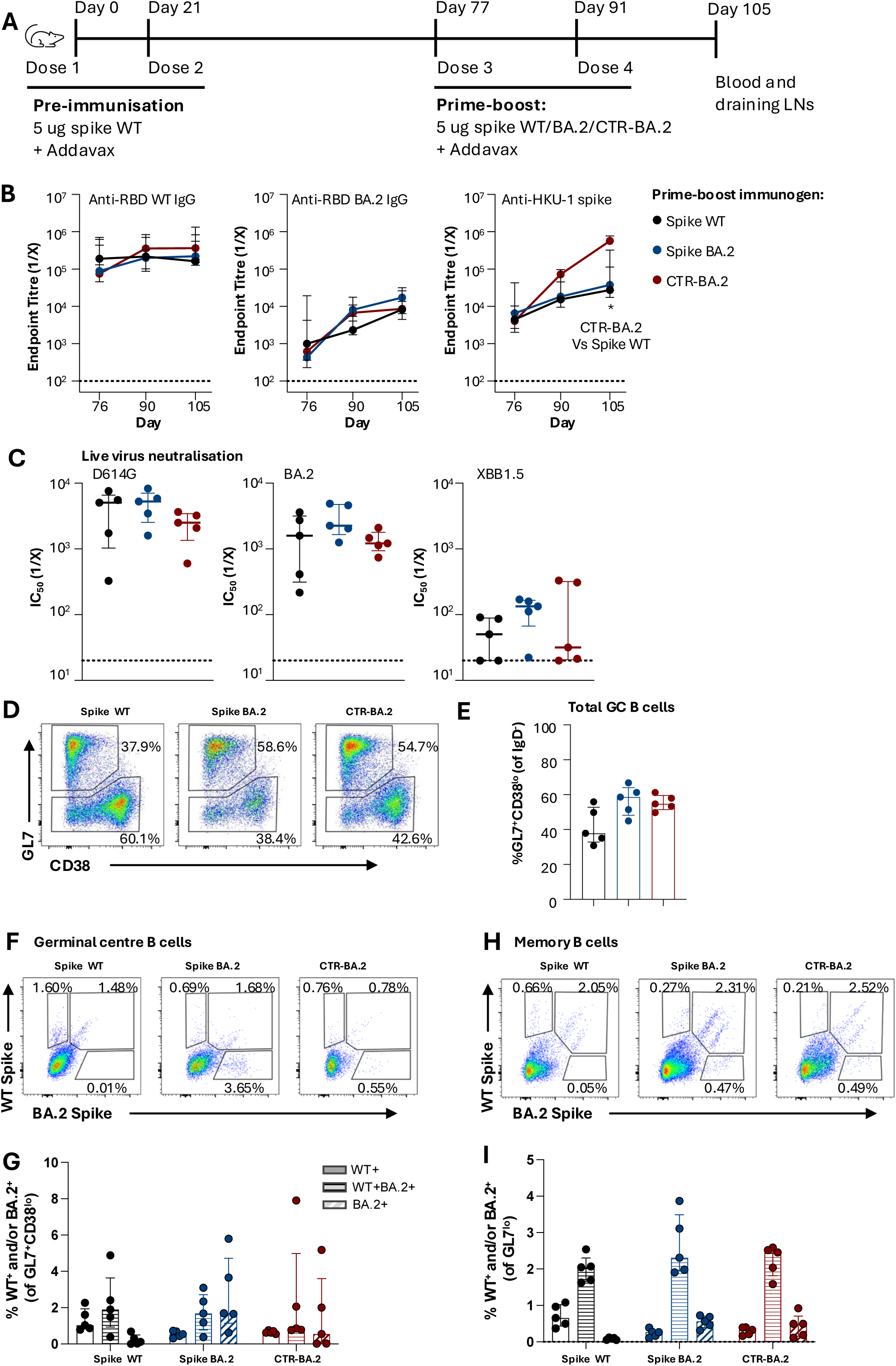
Assessing impact of prior SARS-CoV-2 spike immunity on CTR immunisation in C57BL/6 mice. (A) Immunisation schedule of mice vaccinated first with two doses of 5 ug spike WT Addavax (day 0 and 21) prior to prime-boost immunisation with spike WT, spike BA.2 or CTR BA.2 (day 77 and 91) (n = 5 / group). (B) Longitudinal serum endpoint titres of anti-RBD, -RBD BA.2 and -HKU-1 spike determined by ELISA. * P < 0.05, statistics was assessed by Kruskal-Wallis test. (C) Neutralisation activity in serum at day 105 against SARS-CoV-2 D614G, BA.2 and XBB1.5 assessed by live virus neutralisation assay. (B-C) Dotted lines denote the detection cut off. (D) Representative staining of GC B cells (GL7+ CD38lo) and memory B cells (GL7lo). (E) Frequency of total GC B cells. (F) Representative staining and (G) quantification of WT and/or BA.2 spike-specific GC B cells. (H) Representative staining and (I) quantification of WT and/or BA.2 spike- specific memory B cells.

Previous studies have demonstrated that prior humoral immunity can inhibit the elicitation of secondary GCs upon antigen re-exposure, with serological boosting derived from GC-independent memory B cell recall instead(*28*). We therefore assessed lymph node GC dynamics in CTR-BA.2 vaccinated animals with prior spike WT immunity, finding robust elicitation of GC B cells (Fig 4D-G) and memory B cell (Fig 4H-I and Supplementary Fig S3D) responses comparable to spike WT or BA.2 vaccine groups. In terms of specificity, both GC and memory B cells were predominantly cross-reactive WT^+^ BA.2^+^ populations, with lower frequencies of mono- specific populations (WT^+^ or BA.2^+^) present that matched the antigenicity of the booster immunogens received (Fig 4F-I). Compared to prime-boost vaccinated animals without prior immunity (Fig 2G), CTR-BA.2 vaccination in this pre-immune setting displayed similar frequencies of spike-specific GC B cells 14 days after the last vaccine dose (Supplementary Fig S3E), suggesting that pre-existing antibodies from prior spike WT immunisation did not negatively impact GC elicitation by CTR-BA.2 immunisation. In contrast, significantly higher antigen-specific memory B cells were observed in CTR-vaccinated animals with prior spike exposure, demonstrating recall of spike-specific memory B cells by the CTR-BA.2 (p = 0.0079) (Supplementary Fig S3E). Overall, CTR-BA.2 can efficiently establish de novo GCs and recall memory B cells in animals with pre-existing immunity against SARS-CoV-2 spike WT, while simultaneously boosting neutralising antibody responses against WT or BA.2 to levels comparable to native SARS-CoV-2 spike antigens.

### CTR elicits neutralising antibodies in pigtail macaques with prior human coronavirus immunity

Based on the success of the CTR platform in eliciting neutralising antibodies and recruiting HKU-specific CD4 help in mice, we conducted a prime-boost trial in pigtail macaques (*Macaca nemestrina*), which represent a more genetically diverse pre- clinical model (Fig 5A). As we found no evidence for prior hCoV humoral immunity in macaques upon serological screening (not shown), we first vaccinated ten animals with a cocktail of the 4 common human coronavirus (hCoV) spike proteins (HKU-1 OC43, NL63 and 229E) to mimic the high prevalence of endemic hCoV immunity found in the wider human population. Eight weeks after the second hCoV vaccination, animals were segregated into two groups, receiving either two doses of SARS-CoV-2 spike WT or CTR-WT protein (n = 5/vaccine group). Animals were serially bled post-vaccination for serological assays, with cellular immunity assessed in draining LN collected at necropsy in week 22.

**Figure 5.**
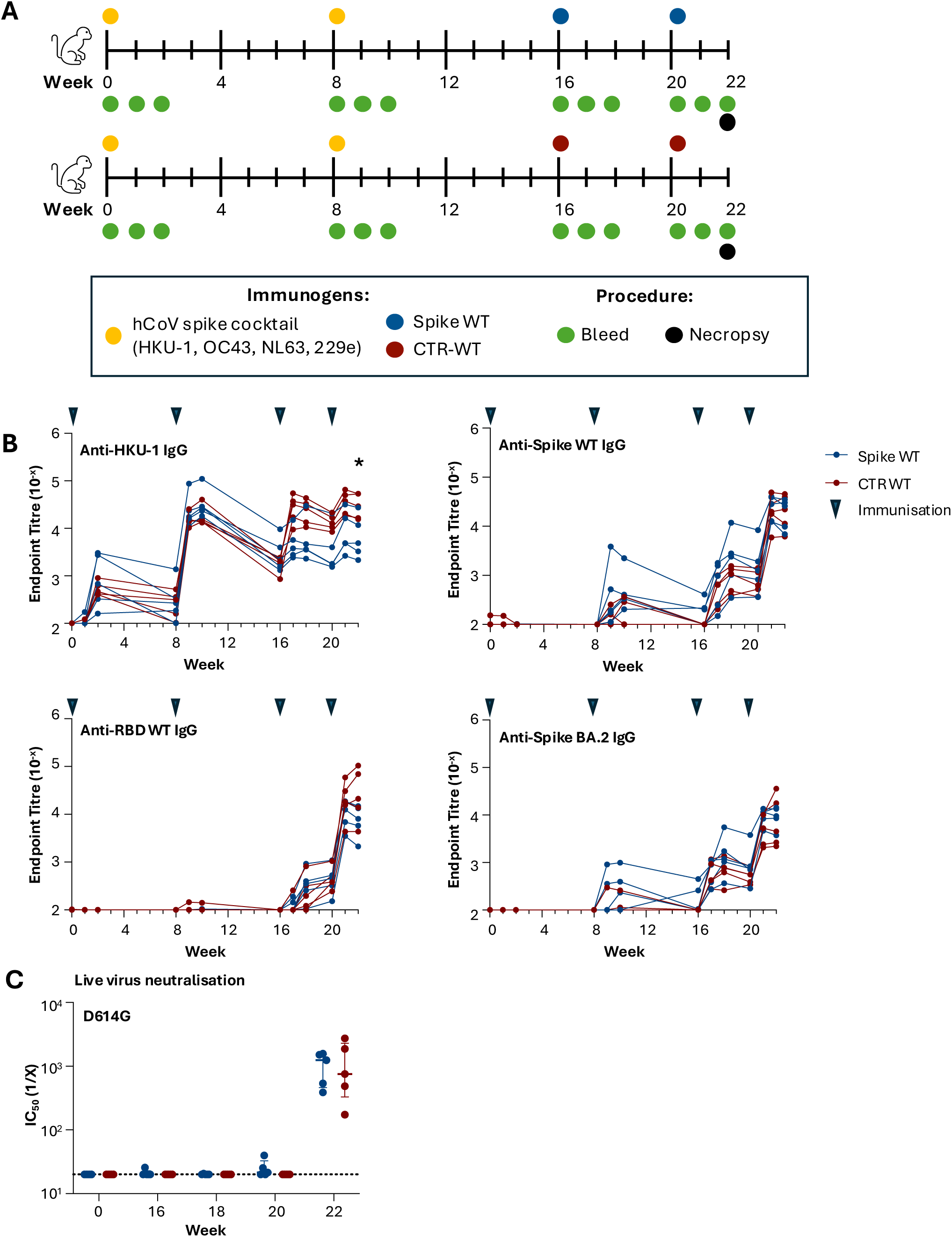
Antibody response and neutralisation kinetics in CTR-WT- and spike WT-vaccinated pigtail macaques. (A) Spike (top) and CTR (bottom) immunisation and bleed schedule in macaques. (n = 5 / group). All animals were immunised with a cocktail of human coronavirus spike proteins (yellow circle; HKU-1 OC43, NL63 and 229e) twice prior to SARS-CoV-2 spike or CTR-WT immunisation. (B) Endpoint titres against HKU-1 spike, SARS-CoV-2 WT spike, SARS-CoV-2 RBD and SARS-CoV-2 BA.2 spike were measured in longitudinal plasma samples by ELISA. Triangles indicate time of immunisations. * P < 0.05, week 22 statistics assessed by Mann-Whitney test. (C) Neutralisation activity against SARS-CoV-2 D614G assessed by live virus neutralisation assay in longitudinal plasma samples. Dotted lines denote the detection cut off.

Serological IgG responses were assessed longitudinally against HKU-1 or SARS-CoV-2 spike antigens (Fig 5B). All animals exhibited seroconversion against HKU-1 spike upon receipt of two doses of the hCoV spike cocktail. Low titres of IgG cross-reactive with spike WT (but not RBD) were detected in all animals post-week 8, likely reflecting responses to cross-reactive S2 epitopes; these responses rapidly waned to low levels (<10^3^). Antibodies to HKU-1 were boosted in macaques immunised with CTR-WT compared to animals boosted with SARS-CoV-2 spike, with significantly elevated titres observed out to necropsy (median: 2.92 x 10^4^ vs 4.88 x 10^3^, respectively; p = 0.0317). Despite CTR-WT carrying only the SARS-CoV-2 RBD, macaques immunised with either CTR-WT or SARS-CoV-2 spike elicited comparable levels of anti-SARS-CoV-2 spike WT IgG titres by week 22 (median: 2.21 x 10^4^ vs 2.58 x 10^4^, respectively). Anti-RBD WT IgG titres appeared higher in three of the five CTR-WT animals compared to the spike WT group (median: 2.12 x 10^4^ vs 8.01 x 10^3^, respectively), although there was substantial variability across the modest numbers of outbred macaques and this did not reach statistical significance. IgG responses against BA.2 spike were elicited at modest and comparable levels between groups (median: 4.50 x 10^3^ vs 9.53 x 10^3^, respectively). Neutralising antibody responses against WT SARS-CoV-2 were only detectable after two doses of spike WT or CTR- WT, and were comparable between vaccine groups (Fig 5C). Overall, CTR efficiently elicits binding and neutralising antibody responses against SARS-CoV-2 in non-human primates with prior hCoV immunity, comparable to levels elicited by canonical spike antigens.

### CTR elicits lymph node TFH and B cell responses in pigtail macaques with prior human coronavirus immunity

GC B cell responses in draining LNs of immunised macaques at week 22 were assessed by flow cytometry using recombinant spike WT, RBD WT and CTR-WT probes (Fig 6A; gating in Supplementary Fig S4A). Significantly larger populations of class-switched GC B cells (p = 0.0078, IgG^+^ IgD^-^ BCL-6^+^ CD95^+^) and probe-specific GC B cells (p = 0.002, CTR-WT^+^ RBD^+^) were identified within ‘draining’ versus ‘non-draining’ LNs (Supplementary Fig S4B), validating the identification and recovery of LNs draining the intramuscular vaccination site. However, the draining LN was not identified in one macaque from the spike WT vaccinated group, which was therefore omitted in subsequent analyses (n = 4 for spike WT group, n = 5 for CTR-WT group) Vaccination with either spike WT or CTR-WT elicited comparable total GC B cell frequencies (median: 22.6% vs 29.9%, respectively) (Fig 6B). Antigen specificity was assessed using two probe combinations: CTR-WT/RBD WT or spike WT/CTR-WT (Fig 6A). All macaques displayed similar frequencies of RBD-specific (CTR-WT^+^ RBD^+^) GC B cells, demonstrating equivalent immunoprominence of the RBD within the HKU-1 scaffold or the native SARS-CoV-2 spike (Fig 6B). As expected, mono-specific populations denoting CoV-2 S2 (CTR-WT^-^, spike WT^+^; 3.25% vs 1.61%) or HKU-1 specific (CTR-WT^+^, spike WT^-^; 4.79% vs 10.8%) GC B cells were found most commonly in spike WT or CTR-WT immunised macaques, respectively. Probe^+^ B cells in the non-GC (BCL-6lo) population, likely demarking the memory pool, were present at lower frequencies within draining LNs and were comparable between vaccine groups (Supplementary Fig S4C).

**Figure 6.**
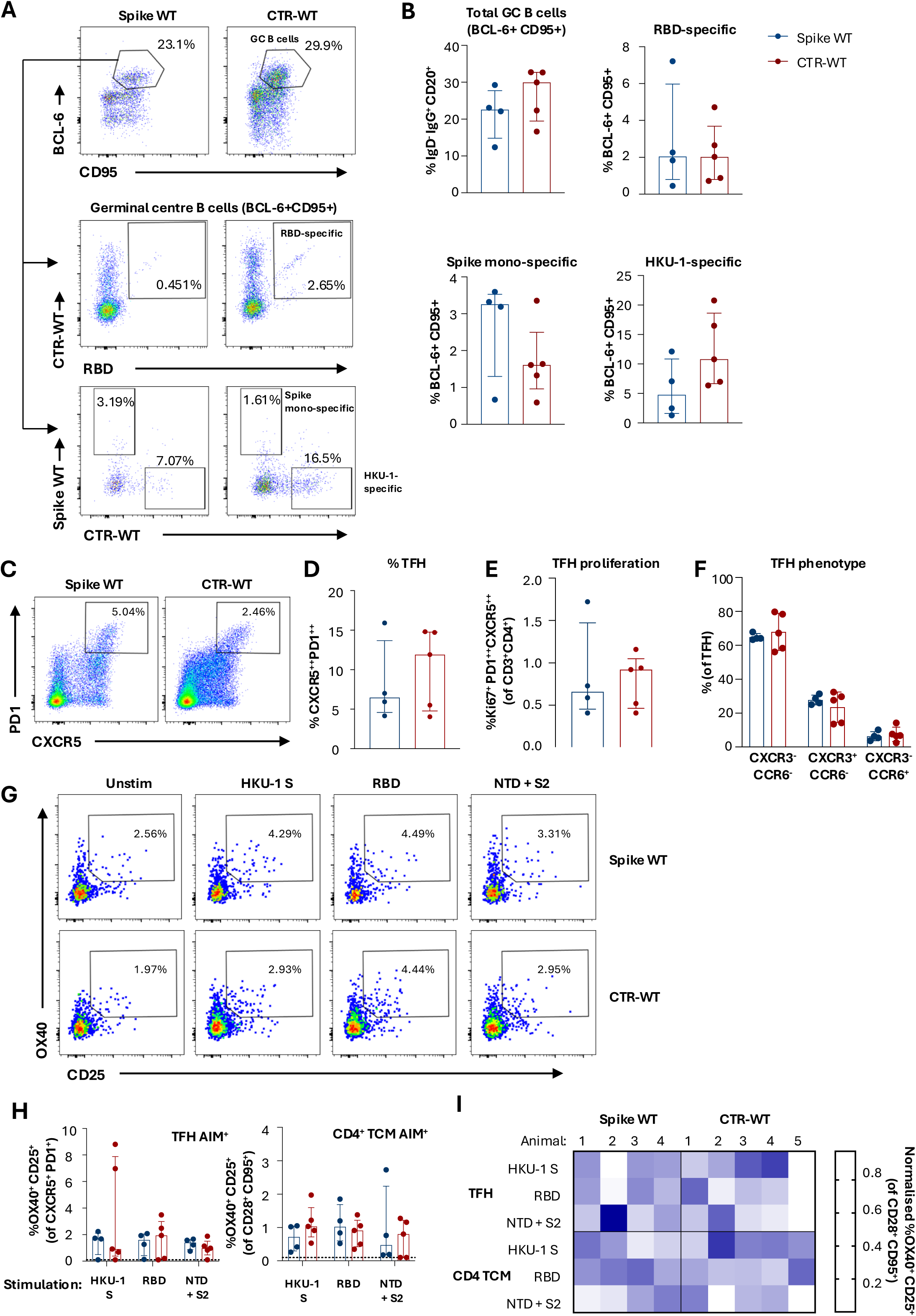
**B and TFH responses in macaques vaccinated with spike WT or CTR- WT**. (A) Representative staining and (B) frequency of SARS-CoV-2 RBD- or spike- specific, and HKU-1 spike-specific GC B cells (CD20+ BCL-6+ CD95+) in vaccine draining lymph nodes of spike WT- (left; n = 4) and CTR-WT-vaccinated (right; n = 5) animals. (C) Representative staining of TFH cells (CD3+ CD4+ CXCR5++ PD-1++). Frequency of (D) total TFH from draining lymph nodes, (E) Ki67+ and (F) CXCR3/CCR6 expression in TFH cells. (G) Representative staining of antigen- specific TFH responses (OX40+ CD25+) in spike WT- (top) and CTR-WT-vaccinated (bottom) animals. (H) Antigen-specific responses in CD4 TFH and TCM (CD3+ CD4+ CXCR5- CD95+ CD28+) subsets following restimulation with HKU-1 spike, SARS- CoV-2 RBD or spike minus RBD (NTD + S2) peptide pools. (I) Heatmap of normalised antigen-specific CD4 TFH and TCM antigen-specific responses frequencies in individual animals.

TFH (CD3^+^ CD4^+^ CXCR5^++^ PD-1^++^) frequencies in draining LN were also assessed following vaccination (Fig 6C; gating in Supplementary Fig S5A). Both vaccines elicited comparable frequencies of bulk TFH (PD-1^++^ CXCR5^++^) and proliferating TFH (Ki67^+^) (Fig 6D-E). While TFH can exhibit variable expression of CXCR3 and CCR6 (potentially reflecting Th1 or Th17-like polarisation) across different infections and vaccine regimens(*29*, *30*), the majority of TFH in the draining LN were CXCR3^-^ CCR6^-^ (Fig 6F).

To assess the antigen specificity of the TFH elicited by each vaccine, we restimulated LN cell suspensions with HKU-1 spike, SARS-CoV-2 RBD, or SARS-CoV-2 NTD+S2 peptide pools in vitro (Fig 6G; gating in Supplementary Fig S5B) and assessed upregulation of CD25 and OX-40(*31*). Median frequencies of AIM^+^ TFH and central memory T cells (TCM, CD28^+^ CD95^+^) were similar between vaccine groups across all specificities (Fig 6H). 8 out of 9 macaques exhibited antigen-specific AIM response above background to at least two of the peptide pools tested, highlighting the diverse sources of CD4 T cell help established by both vaccines and the extent of inter-animal variability in TFH repertoire (Fig 6I). Notably, two of the five CTR-WT animals exhibited particularly high frequencies of HKU-specific TFH, demonstrating the extent to which scaffold-derived T cell epitopes can be recruited into the GC response. These data demonstrate that the immunogenicity of CTR in non-human primates with prior hCoV immunity was comparable to the canonical spike WT antigen, eliciting lymph node GC B and TFH cells of similar phenotypes and specificities.

## Discussion

In this study, we show that the RBD, a key neutralising domain for SARS-CoV-2, can be engineered to be displayed on an hCoV HKU-1 spike glycoprotein backbone. This chimeric antigen displayed a trimeric conformation, with RBD presented in a manner capable of hACE2 binding, and elicited neutralising antibody responses comparable to native spike counterparts in macaque and murine models. This proof-of-concept design highlights a viable strategy for the supplementation of TFH epitopes to augment the intrinsic immunogenicity of the RBD, which we and others previously demonstrated to be relatively poorly immunogenic and ineffective in germinal centre elicitation due to suboptimal CD4 T cell help(*9*, *10*).

Other chimeric spike vaccines comprising domains from up to three coronaviruses have been reported to be immunogenic and elicit neutralising antibodies(*32*, *33*). Similarly, the SARS-CoV spike has been used as a scaffold to display RBDs from Betacoronaviruses for cell entry characterisation(*34*). These studies, together with CTR, suggest there is a high degree of compatibility between domains of coronavirus spike glycoproteins that facilitates the correct display of the RBD. Although this study primarily utilised components of HKU-1 and SARS-CoV-2, which have a high degree of similarity in domain structure and order(*35*, *36*), our modular approach can potentially be extended to use scaffolds derived from other Betacoronaviruses with non-human tropism, such as the bovine or murine hepatitis coronavirus(*37*). Chimeric vaccines with heterologous scaffolds could also be used in serial vaccination regimes to focus RBD neutralising responses via memory recall, akin to approaches used to focus the immune response toward the influenza HA stem domain(*38*, *39*).

Interestingly, despite CTR-WT containing only the RBD of SARS-CoV-2, CTR- immunised macaques elicited total anti-spike and neutralising titres that were equivalent to macaques receiving the native SARS-CoV-2 spike. Spike WT- immunised macaques exhibited 2.6-fold lower anti-RBD titres, with the remaining spike-specific responses likely comprising S2- or NTD-specific antibodies, as previously observed(*9*). Combined with the observation that three of the five CTR- immunised macaques also exhibited the highest anti-RBD titres in the trial, these data highlight the potential potency of RBD in driving neutralising responses. Furthermore, both the macaque and murine models indicated that the HKU-1 scaffold-directed responses did not compete with or constrain the development of RBD-specific antibodies upon CTR vaccination, even in the context of prior HKU-1 immunity. While our CTR design successfully elicited strong RBD-directed responses, the potential for immune responses to be directed towards less or non-neutralising regions (S2 and NTD)(*40*, *41*) in any scaffold design warrants careful consideration to minimise these responses, in particular when applying these design principles to scaffolds from other coronaviruses or from other viral glycoproteins.

Our murine data clearly demonstrate that immunogens with suboptimal CD4 T cell epitopes, such as RBD WT, benefit substantially from covalent linkage to proteins that elicit potent CD4 and TFH responses. As such, the TFH pool in CTR-vaccinated animals was almost exclusively directed toward the HKU-1 scaffold. In contrast, the HLA genetic diversity of non-human primates appears to support much broader recognition of the RBD compared to mice(*9*). Among the ten macaques vaccinated in this study, we observed substantial inter-animal variation in the antigen specificity of LN TFH. Depending on the animal, the dominant population of CD4 help could be derived from the HKU scaffold, the RBD, or NTD/S2 epitopes. Thus, in genetically diverse populations, immunogens that incorporate diverse sources of CD4 T cell epitopes may be key to ensuring broad and uniformly high vaccine immunogenicity.

Overall, our study demonstrates that a chimeric glycoprotein is a viable alternative scaffold structurally comparable to the native SARS-CoV-2 spike glycoprotein. Importantly, this design preserves SARS-CoV-2 RBD presentation in an antigenic conformation enabling elicitation of neutralising responses, while allowing recruitment of scaffold-directed CD4 helper responses to support the humoral response.

## Methods

### Ethics Statement

Animal studies and related experimental procedures were approved by the University of Melbourne Animal Ethics Committee (#22954, #21237) and WEHI Animal Ethics Committee (2020.016). Macaque studies and related experimental procedures were approved by the animal ethics committee at the Monash Animal Research Platform (MARP) (#24539).

### Design and expression of recombinant proteins

Recombinant full length PR8 haemagglutinin (HA) protein with ablated sialic acid binding, ancestral Wuhan Hu-1 (WT) and Omicron BA.2 SARS-CoV-2 spike proteins include C-terminal T4 fibritin foldon, Avitag and his-tag were generated as previously described(*9*, *22*, *42*). Monomeric RBD proteins for WT and BA.2 SARS-CoV-2 including Avitag and his-tag were produced as previously described(*9*, *22*). The HKU-1 spike (NC_006577; AA1-1291) was engineered with two proline stabilisation mutations (S-2P), Avitag and His-tag, as previously described(*21*). CTR-WT was synthesised (GeneArt) with the HKU1 NTD domain (residues 1-348; NC_006577the SARS-CoV-2 RBD (native residues 324-544) and HKU1 S2 ectodomain (residues 622-1282; NC_006577) with 2 proline stabilisation mutations (S-2P), and into two different mammalian expression vectors. To exclude anti-T4 fibritin foldon responses in indirect sandwich ELISA for macaque serology, the C-terminal T4 fibritin folon and Avitag were replaced with GCN4 leucine zipper domain for the following proteins: SARS-CoV-2 spike WT, spike Omicron BA.2 and CTR-WT spike. Proteins were expressed in Expi293 or ExpiCHO cells (Thermo Fisher, Massachusetts USA) using manufacturer’s instructions and purified using Ni-NTA, size exclusion chromatography, and confirmed using SDS-PAGE(*9*).

CTR-BA.2 was similarly designed except with the RBD domain from hCoV- 19/Denmark/DCGC-341558/2022 and without the Avitag. The CTR-BA.2 sequence was codon optimised for expression in CHO cells (Geneart, Thermo Fisher Scientfic), and cloned into the mammalian stable expression vector pXC17.4 (Lonza). CHOK1SV GS-KO® cells (Lonza) were transfected with the linearised vector by electroporation as per the Lonza GS® Xceed Expression System User Manual V8.10 to generate stable cell pools expressing the CTR-BA.2 protein. Pools were selected using methionine sulfoximine. Production of protein from these pools was achieved by an abridged fed batch study as outlined in the Lonza Abridged Fed-Batch Shake-Flask Culture (v9.1) manual. Protein was purified via the His-tag using a HisTrap Excel (Cytiva) affinity column and low pH elution (50mM acetate, 150 mM NaCl, pH 3.6). The eluted protein was adjusted to pH 3.5 with 1 M acetic acid and held for 1 hr for viral inactivation, then neutralized with 1 M Tris to pH 7.35. This was followed by size exclusion chromatography on a Superdex200 HiLoad 26/600 column (Cytiva) in a PBS mobile phase.

### Murine immunisation and euthanasia

5 µg of recombinant SARS-CoV-2 or HKU-1 spike protein were formulated in PBS at 1:1 ratio with Addavax adjuvant (InvivoGen) in total volume of 50 µl. BALB/c or C57BL/6 mice aged 6-12 weeks were anaesthetised to required depth (i.e. hind legs do not move in response to light pressure) by isoflurane inhalation prior to intramuscular vaccination, with oxygen flow at 2 L/min and isofluorane vaporiser set to 3 (Stinger Anaesthetic Machine). Once anaesthetised, animals are immunised with 50 µL of vaccine to the injection site using a 29G needle (left quadriceps muscle or right deltoid muscle). Prior to tissue collection, mice were killed using CO2 asphyxiation using a delivery of 50% of chamber volume per min, followed by secondary method of killing using cervical dislocation.

### Macaque immunisation and euthanasia

Pigtail macaques (*Macaca nemestrina*) were housed in the Monash Animal Research Platform. For vaccinations, macaques were restrained in a primate housing unit with a squeeze-back mechanism and performed without anaesthesia while the animal was conscious. 10 male animals, aged 2-4 years, were vaccinated with two doses of 40 μg of hCoV spike cocktail (HKU-1, OC43, 229E and NL63) and was followed by two doses of 25 µg CTR-WT or SARS-CoV-2 WT spike. All immunisations were formulated with Monophosphoryl Lipid A (MPLA) liposomes (Polymun) administered intramuscularly in the left and right quadriceps. For an unrelated study, animals were concurrently vaccinated in the left and right deltoids with HIV-trimeric envelope protein gp140 (SOSIP) immunogens (50 μg) formulated with MPLA and 1% tattoo ink. For euthanasia, macaques were restrained in a primate housing unit with a squeeze-back mechanism. Animals were then administered an intramuscular injection of sedative cocktail containing Medetomidine, Butorphanol, Atropine and Ketamine using a 22G needle followed by Sevoflurane by inhalation. Once anaesthetised, animals were euthanised with an overdose of Pentobarbitone via intravenous injection with a 23G needle in the saphenous vein. Macaques were necropsied 14 days after the last immunisation.

### Negative-staining electron microscopy

CTR-BA.2 spike was diluted in 20 mM HEPES pH 7.5, 150 mM NaCl to 0.1 mg/ml. Carbon coated 400 micron mesh copper TEM grids were glow discharged for 1 min at a current of 15 mA and 4 μl of the sample was then allowed to adsorb for 1 min. Grids were then stained using 4 μl 1% Uranyl acetate for 1 min and blotted dry. Micrographs were collected on a Tecnai G2 Spirit TEM microscope (FEI) running at 120 keV equipped with an Eagle 4x4k CMOS camera at a magnification of 52,000 (pixel size 4.15 Å). Particle picking and reference-free 2D classification were performed in CryoSPARC 4.6.2(*43*).

### SARS-CoV-2 D614G N501Y virus challenge and measurement of lung viral burden

SARS-CoV-2 challenge and lung viral load measurement were performed as previously described(*44*). Briefly, live SARS-CoV-2 were conducted in an Office of Gene Technology Regulator-approved Physical Containment Level 3 facility at the Walter and Eliza Hall Institute of Medical Research (Cert-3621). All animal procedures were approved by the Walter and Eliza Hall Institute of Medical Research Animal Ethics Committee (2020.016). BALB/c mice were infected with SARS-CoV-2 clinical isolate hCoV-19/Australia/VIC2089/2020, which harbours the naturally occurring D614G and N501Y mutations; N501Y enables infection of wild-type mice via the murine ACE2 receptor. Infection was performed using an inhalation exposure system (Glas-Col, LLC) for 45 min loaded with 1.5 × 10^7^ SARS-CoV-2 TCID50. Animals were humanely euthanised and lungs removed and homogenised in a Bullet Blender (Next Advance Inc) in 1 mL DME media (ThermoFisher) containing steel homogenisation beads (Next Advance Inc). Samples were clarified by centrifugation at 10,000 x g for 5 mins before virus quantification by TCID50 assays. SARS-CoV-2 live virus quantification by TCID50 was determined by plating 1:7 serially-diluted lung tissue homogenate onto confluent layers of Vero cells (clone CCL81) in DME media (ThermoFisher) containing 0.5 μg/mL trypsin-TPCK (ThermoFisher) in replicates of six on 96-well plates. Plates were incubated at 37 °C supplied with 5% CO2 for four days before measuring cytopathic effect under phase contrast light microscope. The TCID50 calculation was performed using the Spearman and Kärber method(*45*).

### ELISA for assessment of serological responses and antigenicity

For assessment of murine serology, 96-well Maxisorp plates (Thermo Fisher Scientific) were coated overnight at 4°C with recombinant RBD WT, RBD BA.2, or HKU-1 S (2 µg/mL). After blocking with 5% skim milk powder in PBS for 1 h, serially diluted mouse antisera were added and incubated for 2 h at room temperature. Bound antibody was detected using a 1:15,000 dilution of peroxidase-labelled goat anti-mouse IgG (Seracare). Plates were developed using 3,3’,5,5’ Tetramethylbenzidine (TMB) substrate (Invitrogen), stopped with sulphuric acid, and read at 450 nm. Endpoint titres were calculated using GraphPad Prism as the reciprocal serum dilution giving signal 2× background using a fitted curve (four-parameter log regression).

For macaque serology, 96-well Maxisorp plates (Thermo Fisher Scientific) were coated with 1 µg/mL anti-his tag antibody (Genescript) overnight at 4°C. After blocking with 5% skim milk powder in PBS, 2 µg/mL recombinant WT spike, BA.2 spike, RBD WT, or HKU-1 spike was added and incubated for 1 h at room temperature. Duplicate wells of serially diluted plasma were added and incubated for 2 h at room temperature. Plates were washed prior to the incubation of HRP-conjugated secondary antibody for macaques (1:8000; anti-monkey IgG; Southern Biotech) for 1 h at room temperature. Plates were washed before developing with TMB substrate (Invitrogen) and read at 450nm. Endpoint titres were calculated as the reciprocal plasma dilution equal to 2x background using a fitted curve (four parameter log regression).

For epitope mapping, 96-well Maxisorp plates (Thermo Fisher Scientific) were coated overnight at 4°C with recombinant WT spike, BA.2 spike, RBD WT, RBD BA.2, CTR- WT, CTR-BA.2, or HKU-1 spike (2 µg/mL). Blocking and dilution buffers used for this assay were PBS and 1% FCS. Monoclonal antibodies (mAbs) specific for SARS-CoV- 2 RBD, NTD, S2 and negative control PGT121 were isolated and synthesised, as previously described(*46*). Serially diluted monoclonal antibodies were added with a starting concentration of 10 µg/mL, followed by a 2 h incubation at room temperature. Antibody binding was detected using a 1:20,000 dilution of horseradish peroxidase (HRP)-conjugated rabbit anti-human IgG (Dako). Plates were washed before developing with TMB substrate (Invitrogen) and read at 450nm.

### hACE2 binding assay

To detect human ACE2 (hACE2) binding, 96-well plates were coated overnight at 4°C with recombinant S WT, S BA.2, RBD WT, RBD BA.2, CTR-WT, CTR-BA.2, or HKU-1 S (2 µg/mL) in duplicates. The plates were blocked with PBS and 1% BSA for 1 h, and biotinylated hACE2 was added, followed by a 1 h incubation at room temperature. Binding activity was detected using Pierce™ High Sensitivity Streptavidin-HRP (Thermo Fisher Scientific). Plates were washed before developing with TMB substrate (Invitrogen) and read at 450nm.

### Flow cytometric detection of B cells

For use as flow cytometry probes, spike, HA and RBD proteins were biotinylated using BirA (Avidity) and labelled with the sequential addition of streptavidin (SA) conjugated to BV711 (BD), BV510 (BD), PE (ThermoFisher) or APC (ThermoFisher), as previously described(*9*). Alternatively, CTR-WT spike with GCN4 leucine zipper domain was directly labelled using PE and APC Conjugation Lightning-Link Kit (Abcam) following the manufacturer’s protocol for use as probes.

For murine studies, lymph nodes were harvested into RF10 media (RPMI 1640, 10% FCS, 1x penicillin-streptomycin-glutamine: Life Technologies, Thermo Fisher Scientific). Single cell suspensions were prepared by mechanical dissociation through a 70 µM cell strainer. Cells were stained with Aqua viability dye (Thermo Fisher) and Fc-blocked with anti-CD16/32 antibody (clone 93, BioLegend). Cells were then labelled with PE- and APC-conjugated spike probes, and the following antibodies at 4°C: F4/80 BV786 (BM8; BioLegend; 1:150), CD3 BV786 (145-2C11; BioLegend; 1:750), strepavidin BV786 (BD), CD45 APC-Cy7 (30-F11; BD; 1:300), CD38 PeCy7 (90; BioLegend; 1:750), GL7 AF488 (GL7; BioLegend; 1:300), IgD BUV395 (11-26c.2a; BD; 1:300), and B220 BUV737 (RA3-6B2; BD; 1:300). Cells were washed twice with PBS + 1% FCS and fixed using 1% formaldehyde (Polysciences) and acquired on a BD LSR Fortessa using BD FACS Diva.

For macaque studies, draining iliac lymph nodes were harvested and passed through a 70um filter. Single cell suspensions were washed in RPMI, cryopreserved in 90% FCS/10% DMSO and stored in liquid nitrogen until use. Cryopreserved lymph node suspensions were thawed and split into two aliquots to be stained with two combinations of probes: HA PR8 BV510, RBD BV711, RBD APC and CTR-WT PE or HA PR8 BV510, CTR-WT APC, CTR-WT PE and spike WT BV711. Prior to surface stain, cells were stained with Aqua viability dye (Invitrogen). Along with the different probe combinations, macaque cells were stained with the following antibody cocktail for 30 mins at 4°C: CD14 BV510 (M5E2; BL; 1:300), CD3 BV510 (OKT3; Biolegend; 1:100), CD8a BV510 (RPA-T8; Biolegend; 1:400), CD16 (3G8; Biolegend; 1:500), CD10 BV510 (HI10a; Biolegend; 1:75), Streptavidin BV510 (BD), IgG BV786 (G18- 145; BD; 1:75), CD20 APC-Cy7 (2H7; Biolegend; 1:150), IgD AlexaFluor488 (Polyclonal; Southern Biotech; 1:150) and CD95 BUV395 (DX2; BD; 1:200). Macaque cells were permeabilised and fixed for 45 mins in the dark at 4°C using BD Pharmingen Transcription Buffer Set and transcriptionally stained with BCL-6 PE-Cy7 (G18-145; BD; 1:12.5) for 40 mins at room temperature. Cells were then fixed in 1% formaldehyde and acquired on a BD LSR Fortessa using BD FACS Diva.

### Flow cytometric detection of ex vivo and antigen-specific TFH

To identify antigen-specific TFH in mice, LN single cell suspensions were cultured for 18 h at 37°C in RF10 supplemented with 20 μM A438079 to inhibit TFH cell death. For macaques, cryopreserved LN suspensions were thawed, rested for 2 h, and stimulated in RF10 for 20 h, as previously described(*9*). Cells were stimulated with overlapping peptide pools spanning the HKU-1 spike protein (PepMix, JPT Peptide Technologies), the WT RBD (custom peptide array, Genscript) or the WT spike protein without RBD (custom peptide array, Genscript) at 2μg/mL/peptide. In some experiments, stimulations were performed with recombinant RBD BA.2, RBD WT, spike BA.2, spike WT or HKU-1 spike proteins at 2 μg/mL. Anti-CD154 BV650 (MR1; Biolegend; for mice) or PE (TRAP-1; BD; for macaques) was added to the culture for the full incubation. As a negative control, cells were cultured with an equivalent volume of DMSO (for peptide stimulations) or PBS (for protein stimulations).

After culture, cells were washed in PBS and stained with live/dead viability dye (Life Technologies) for 3 mins at room temperature. Mouse cells were then stained with the following antibody cocktail for 30 mins at 4°C in the dark: CD3 APC-Fire750 (145- 2C11; Biolegend; 1:50), CD25 BB515 (PD61; BD; 1:50), PD-1 BV786 (29F.1A12; Biolegend; 1:100), CXCR5 BV421 (L138D7; Biolegend; 1:50), CD4 BUV737 (RM4-5; BD; 1:200), OX-40 PeCy7 (OX-86; Biolegend; 1:100), B220 BV605 (RA3-6B2; BD; 1:100), CD44 AlexaFluor700 (IM7; BD; 1:100) and F4/80 PE-Dazzle 594 (T45-2342; BD; 1:100). Macaque cells were stained with the following antibody cocktail under the same conditions: CD20 BV510 (2H7; BD; 1:100), CD3 AlexaFluor700 (SP34-2; BD; 1:100), CD4 BV605 (L200; BD; 1:100), PD-1 BV421 (EH12.27H; Biolegend; 1:50), CD8 BV650 (RPA-T8; Biolegend; 1:400), CD28 BV711 (CD28.2; BD; 1:25), CD95 BUV737 (DX2; BD; 1:200), CXCR3 PE Dazzle 594 (G02H57; Biolegend; 1:50), CCR6 BV785 (G034E3; Biolegend; 1:100), CXCR5 PECy7 (MU5UBEE; ThermoFisher; 1:33), OX-40 BUV395 (L106; BD; 1:33), CXCR4 APC (12G5; BD; 1:100) and CD25 BB515 (M-A251; BD; 1:50). Cells were then fixed in 1% paraformaldehyde and acquired on a BD LSR Fortessa.

To identify different T cell phenotypes in macaques, cryopreserved LN suspensions were thawed and stained with live/dead viability dye (Life Technologies) for 3 mins at room temperature. Macaque cells were next stained with the following antibody cocktail away from light at 4°C for 30 mins: ICOS PerCPCy5.5 (C398.4A, Biolegend; 1:50), CD3 AlexaFluor700 (SP34-2; BD; 1:100), PD-1 BV421 (EH12.27H; Biolegend; 1:50), CD20 BV510 (2H7; BD; 1:100), CD4 BV605 (L200; BD; 1:100), CD8 BV650 (RPA-T8; Biolegend; 1:400), CCR6 BV785 (G034E3; Biolegend; 1:100), CXCR3 PE Dazzle 594 (G02H57; Biolegend; 1:50), CXCR5 PECy7 (MU5UBEE; ThermoFisher; 1:33) and CD95 BUV737 (DX2; BD; 1:150). Macaque cells were permeabilised and fixed for 45 mins in the dark at 4°C using BD Pharmingen Transcription Buffer Set and transcriptionally stained with BCL6 AlexaFluor647 (IG191E/A8; Biolegend; 1:12.5) and Ki67 BUV395 (B56; BD; 1:20) for 40 mins at room temperature. Cells were then fixed in 1% paraformaldehyde and acquired on a BD LSR Fortessa.

### SARS-CoV-2 microneutralisation assay

A replicating SARS-CoV-2 microneutralisation assay was performed as previously described(*47*), against the ancestral variant with D614G mutation, Omicron BA.2 and Omicron XBB.1.5 isolates. Infectivity of virus stocks was then determined by titration on HAT-24 cells (a clone of transduced HEK293T cells stably expressing human ACE2 and TMPRSS2)(*48*). Virus stocks were titrated in quintuplicate in three independent experiments to obtain mean 50% infectious dose (ID50) values.

To determine serum neutralisation activity, heat-inactivated mouse serum samples were diluted 3-fold (1:20–1:43,740) in duplicate and incubated with SARS-CoV-2 virus at a final concentration of 2 × ID50 at 37°C for 1 h. Next, 40,000 freshly trypsinised HAT-24 cells in DMEM with 5 % FCS were added and incubated at 37°C. “Cells only” and “Virus+Cells” controls were included to represent 0% and 100% infectivity respectively. After 48 h, 10 μL of alamarBlue Cell Viability Reagent (ThermoFisher) was added into each well and incubated at 37 °C for 1 h. The reaction was then stopped with 1% SDS and read on a FLUOstar Omega plate reader. The relative fluorescent units (RFU) measured were used to calculate %neutralisation with the following formula: (“Sample” - “Virus+Cells”) ÷ (“Cells only” - “Virus+Cells”) × 100. IC50 values were determined using four-parameter non-linear regression in GraphPad Prism with curve fit constrained to have a minimum of 0% and a maximum of 100% neutralisation.

### Statistics

Data is generally presented as median +/− interquartile range. Statistical significance was assessed by Mann–Whitney or Wilcoxon tests. Curve fitting was performed using four parameter logistic regression. All statistical analyses were performed using Prism (GraphPad). Flow cytometry data was analysed in v10.

## Acknowledgements

The authors acknowledge the facilities and the scientific and technical assistance of the Bioresources Facility (BRF) and Melbourne Cytometry Platform, at the University of Melbourne. Human ACE2 assay reagents were generously provided by Bruce D Wines, P Mark Hogarth, at the Burnet Institute, Melbourne, Australia. The authors acknowledge Anne Gibbon and the Monash Animal Research Platform (Monash University) for their assistance with the non-human primate trials. The authors also acknowledge the facilities and the scientific assistance of the National Biologics Facility (NBF) at The University of Queensland. NBF is supported by Therapeutic Innovation Australia (TIA). TIA is supported by the Australian Government through the National Collaborative Research Infrastructure Strategy (NCRIS) program.

## Author contributions

Conceptualization: SJK, AKW, JJ, HXT

Formal Analysis: VZ, WSL, LM, PP, AKW, JJ, HXT Funding Acquisition: SJK, AKW, JJ, HXT

Investigation: VZ, WSL, LM, LB, PP, IBA, JPC, KCD, MD, AKW, JJ, HXT Methodology: VZ, RE, AK, HGK, CEM, MG, KH, MLJ, BH, AKW, JJ, HXT Resources: MLJ, MP, WHT, BH, SJK, AKW, JJ, HXT

Supervision: AKW, JJ, HXT

Writing – original draft: VZ, AKW, JJ, HXT

Writing – review & editing: VZ, WSL, SJK, AKW, JJ, HXT

## Supplementary Figures

**Figure S1.**
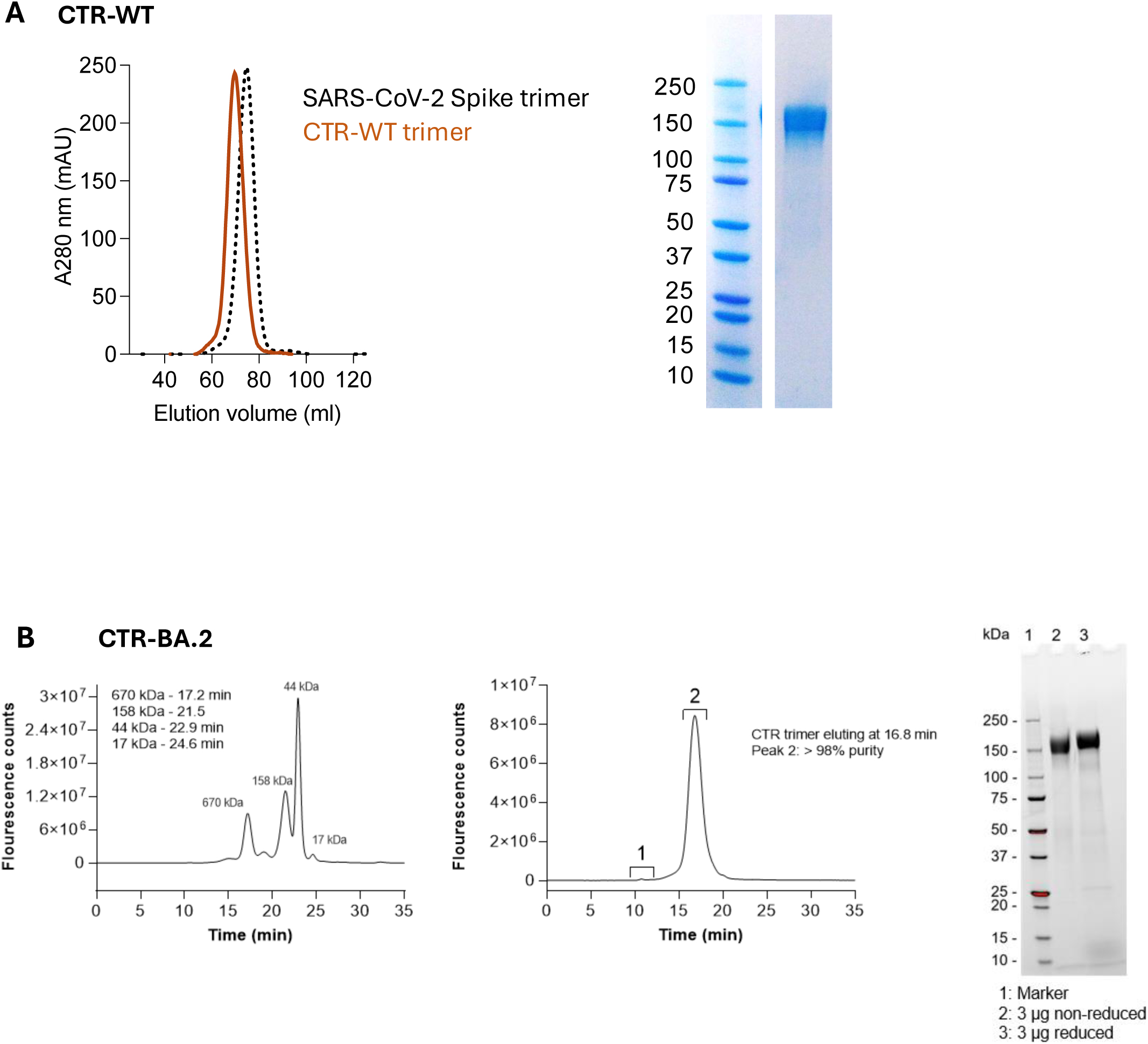
**Validation of purified chimeric trimer RBD glycoproteins**. Affinity/size exclusion chromatography trace and expected size shown by SDS-PAGE for **(A)** CTR-WT and **(B)** CTR-BA.2 glycoproteins.

**Figure S2.**
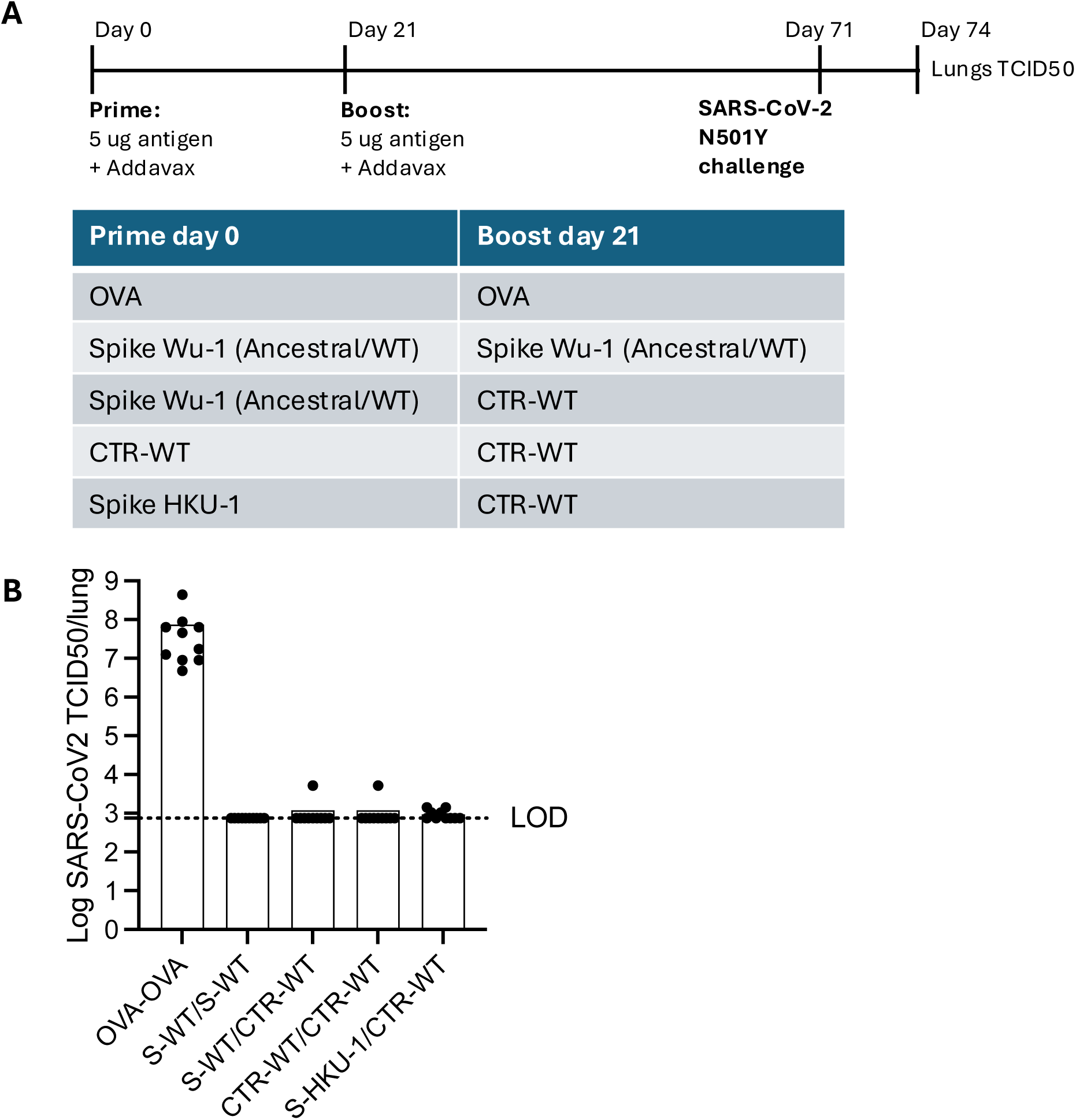
**Immune challenge of BALB/c mice with mouse infectious SARS-CoV-2 virus**. **(A)** Immunisation schedule and challenge with SARS-CoV-2 N501Y virus. **(B)** Viral load measurement of SARS-CoV-2 N501Y virus in lungs of mice three days post-challenge. Dotted lines denote limit of detection.

**Figure S3.**
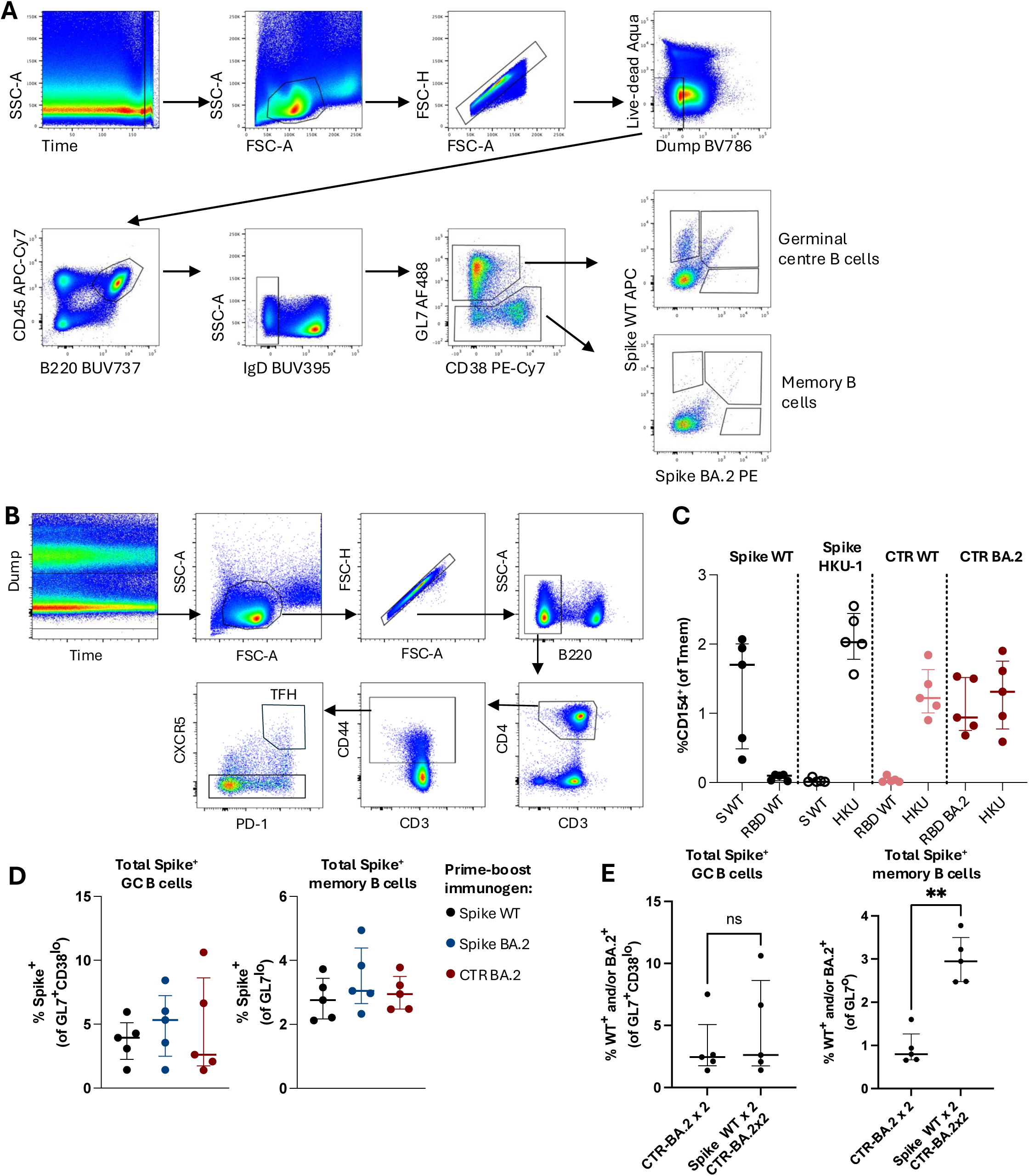
Identification of B cells and CD4 T cells in mice. **(A)** B cells were identified by SSC-A vs time, FSC-A vs SSC-A, followed by doublet exclusion (FSC-A vs FSC-H). Live and CD3^-^ F4/80^-^ streptavidin^-^ (dump channel) cells were gated and CD45^+^ B220^+^ IgD^-^ B cells identified. Germinal centre (GL7^+^ CD38^lo^) or memory (GL7^lo^) B cells were then assessed for binding to SARS-CoV-2 spike WT and/or BA.2 probes. **(B)** Gating strategy for lymph node CD4 TFH or bulk Tmem cells, based on CD44^hi^, PD-1 and CXCR5 expression. **(C)** Frequencies of antigen-specific CD154^+^ bulk Tmem cells upon restimulation with whole spike or RBD proteins. **(D)** Frequencies of total spike-specific GC and memory B cells at day 105 in mice after vaccination schedule in Figure 4A. **(E)** Comparison of total spike-specific GC and memory B cells in mice with or without prior spike WT immunity 14 days after last immunisation. ** P < 0.01, statistics assessed by Mann-Whitney test.

**Figure S4.**
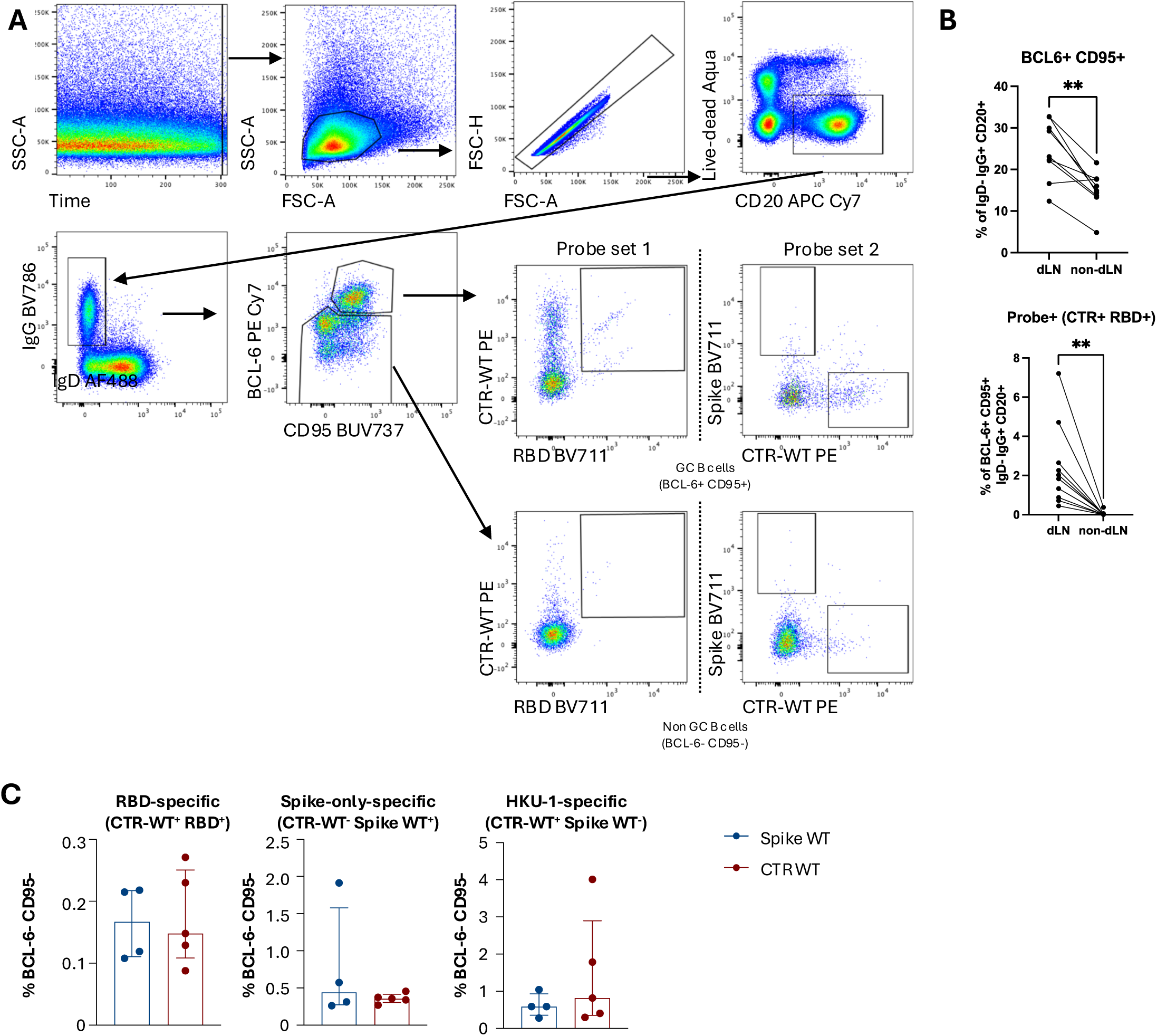
Identification of lymph node B cells in macaques. **(A)** B cells were first identified by SSC-A vs Time, FSC-A vs SSC-A gating, followed by doublet exclusion (FSC-A vs FSC-H). Cells were then gated on dump^-^ (CD3^-^ CD8^-^ CD14^-^ CD10^-^ CD16^-^ streptavidin^-^) live CD20^+^ B cells. GC and non-GC B cells were identified from class-switched IgD^-^ B cells with BCL-6^+^ CD95^++^ or BCL-6^lo^ staining, respectively. Antigen-specific cells were identified using two probe combinations: CTR-WT/RBD WT or spike WT/CTR-WT. **(B)** Comparison of total and antigen-specific GC B cell frequencies between paired draining and non-draining lymph nodes of vaccinated macaques. ** P < 0.01, statistics assessed by Wilcoxon test. **(C)** Frequency of antigen-specific non-GC B cells in draining lymph nodes.

**Figure S5.**
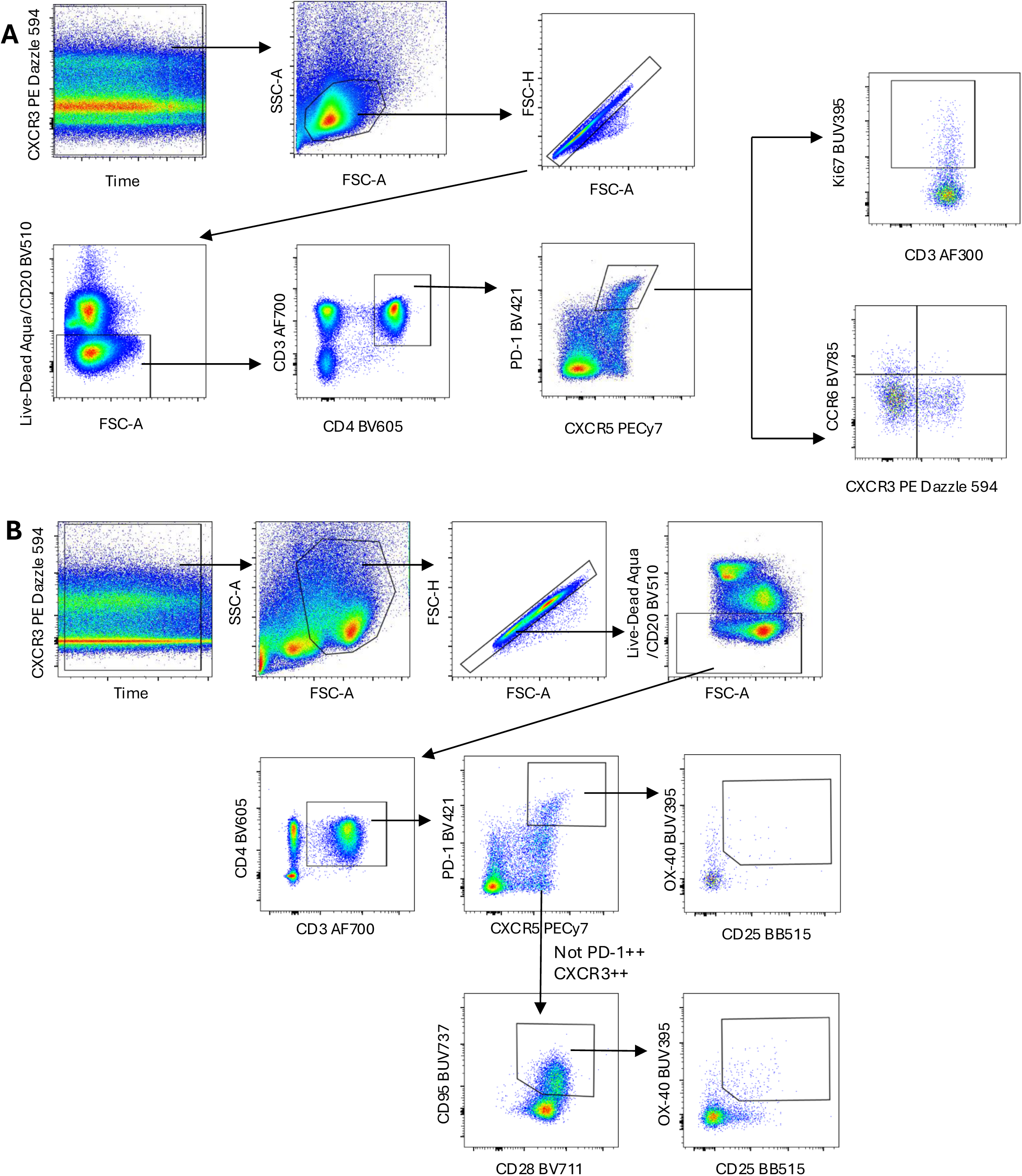
Identification of lymph node CD4 T cells in macaques. **(A)** TFH cells were first identified by Time, FSC-A vs SSC-A gating, followed by doublet exclusion (FSC-A vs FSC-H). Cells were then gated on CD20- live cells, followed by CD3^+^ and CD4^+^ expression. Bulk TFH cells were identified as PD-1^++^ CXCR5^++^, with proliferating TFH gated through Ki67^+^. Th1 or Th17-like polarisation was further identified by CXCR3 and CCR6 expression. **(B)** Identification of antigen-specificity of TFH (PD-1^++^ CXCR5^++^) and TCM (CD28^+^ CD95^+^) subsets with similar upstream gating and by downstream gating of CD25 and OX-40 upregulation.

